# A large-scale, standardized physiological survey reveals higher order coding throughout the mouse visual cortex

**DOI:** 10.1101/359513

**Authors:** Saskia E. J. de Vries, Jerome Lecoq, Michael A. Buice, Peter A. Groblewski, Gabriel K. Ocker, Michael Oliver, David Feng, Nicholas Cain, Peter Ledochowitsch, Daniel Millman, Kate Roll, Marina Garrett, Tom Keenan, Leonard Kuan, Stefan Mihalas, Shawn Olsen, Carol Thompson, Wayne Wakeman, Jack Waters, Derric Williams, Chris Barber, Nathan Berbesque, Brandon Blanchard, Nicholas Bowles, Shiella Caldejon, Linzy Casal, Andrew Cho, Sissy Cross, Chinh Dang, Tim Dolbeare, Melise Edwards, John Galbraith, Nathalie Gaudreault, Fiona Griffin, Perry Hargrave, Robert Howard, Lawrence Huang, Sean Jewell, Nika Keller, Ulf Knoblich, Josh Larkin, Rachael Larsen, Chris Lau, Eric Lee, Felix Lee, Arielle Leon, Lu Li, Fuhui Long, Jennifer Luviano, Kyla Mace, Thuyanh Nguyen, Jed Perkins, Miranda Robertson, Sam Seid, Eric Shea-Brown, Jianghong Shi, Nathan Sjoquist, Cliff Slaughterbeck, David Sullivan, Ryan Valenza, Casey White, Ali Williford, Daniela Witten, Jun Zhuang, Hongkui Zeng, Colin Farrell, Lydia Ng, Amy Bernard, John W. Phillips, R. Clay Reid, Christof Koch

## Abstract

To understand how the brain processes sensory information to guide behavior, we must know how stimulus representations are transformed throughout the visual cortex. Here we report an open, large-scale physiological survey of neural activity in the awake mouse visual cortex: the Allen Brain Observatory Visual Coding dataset. This publicly available dataset includes cortical activity from nearly 60,000 neurons collected from 6 visual areas, 4 layers, and 12 transgenic mouse lines from 221 adult mice, in response to a systematic set of visual stimuli. Using this dataset, we reveal functional differences across these dimensions and show that visual cortical responses are sparse but correlated. Surprisingly, responses to different stimuli are largely independent, e.g. whether a neuron responds to natural scenes provides no information about whether it responds to natural movies or to gratings. We show that these phenomena cannot be explained by standard local filter-based models, but are consistent with multi-layer hierarchical computation, as found in deeper layers of standard convolutional neural networks.

## Introduction

Traditional understanding, based on several decades of research, suggests that neurons early in the visual pathway are broadly responsive and become more selective and specialized through a series of hierarchical processing stages^1–4^. However, the computations and mechanisms required for such transformations remain unclear. A key challenge results from the fact that our understanding of the mammalian visual system is the result of many small studies, recording responses from different stages in the circuit, using different stimuli and different analyses.^5^ The inherent experimental selection biases and lack of standardization of this approach introduce additional obstacles to creating a cohesive understanding of cortical function. To address these differences, we conducted a survey of visual responses across multiple layers and areas in the mouse visual cortex, using a diverse set of visual stimuli. This survey was executed in pipeline fashion, with standardized equipment and protocols and with strict quality control measures not dependent upon stimulus-driven activity (**see Methods, Supplemental Figures 1-8**).

Previous work in mouse has revealed functional differences among cortical areas in layer 2/3 in terms of the spatial and temporal frequency tuning of the neurons in each area.^6,7^ However, it is not clear how these differences extend across layers and across diverse neuron populations. Here we extend such functional studies to include 12 Cre-defined neuron populations, including excitatory populations across 4 cortical layers (from layer 2/3 to layer 6), and two inhibitory populations (Vip and Sst). Further, it is known that stimulus statistics affect visual responses, such that responses to natural scenes cannot be well predicted by responses to noise or grating stimuli^8–11^. To examine the extent of this discrepancy in the mouse visual cortex, and whether it varied across areas and layers, we designed a stimulus set that included both artificial (gratings and noise) and natural (scenes and movies) stimuli. While artificial stimuli can be easily parameterized and interpreted, natural stimuli are likely to be closer to what is ethologically relevant to the mouse. Finally, as recording modalities have enabled recordings of larger and larger populations of neurons, it has become clear that populations might code visual and behavioral activity in a way that is not apparent by considering single neurons alone.^12^ Here we imaged populations of neurons (mean 118 ± 82, st. dev, for excitatory populations) to explore both single neuron and population coding properties.

We find that 74% of neurons in the mouse visual cortex respond to at least one of these visual stimuli, many showing classical tuning properties, such as orientation and direction selective responses to gratings. These tuning properties reveal functional differences across cortical areas and Cre lines. The responses to all stimuli are highly sparse, both in terms of lifetime and population sparseness. We demonstrate that for all cells the visual responses are better fit by a quadratic “complex cell” model than by a linear-nonlinear “simple cell” model. Importantly, we find that the responsiveness to different stimuli is largely independent, i.e. cells that respond to natural movies do not necessarily respond to natural scenes. These properties are not consistent with a traditional Gabor-style spatio-temporal wavelet basis, but rather are to be expected in deeper layers of a multi-layer hierarchical network.

## Results

Using adult C57BL/6 mice (mean age 108 ± 17 days st. dev) that expressed a genetically encoded Ca2+ sensor (GCaMP6f) under the control of specific Cre-line drivers (10 excitatory lines, 2 inhibitory lines, **Supplemental Figure 7**), we imaged the activity of neurons in response to a battery of diverse visual stimuli. Data was collected from 6 different cortical visual areas (V1, LM, AL, PM, AM, and RL) and 4 different cortical layers. Visual responses of neurons at the retinotopic center of gaze were recorded in response to drifting gratings, flashed static gratings, locally sparse noise, natural scenes and natural movies (**Figure 1f**), while the mouse was awake and free to run on a rotating disc. In total, 59,526 neurons were imaged from 410 experiments, each consisting of three one-hour imaging sessions (**Table 1**).

**Figure 1:**
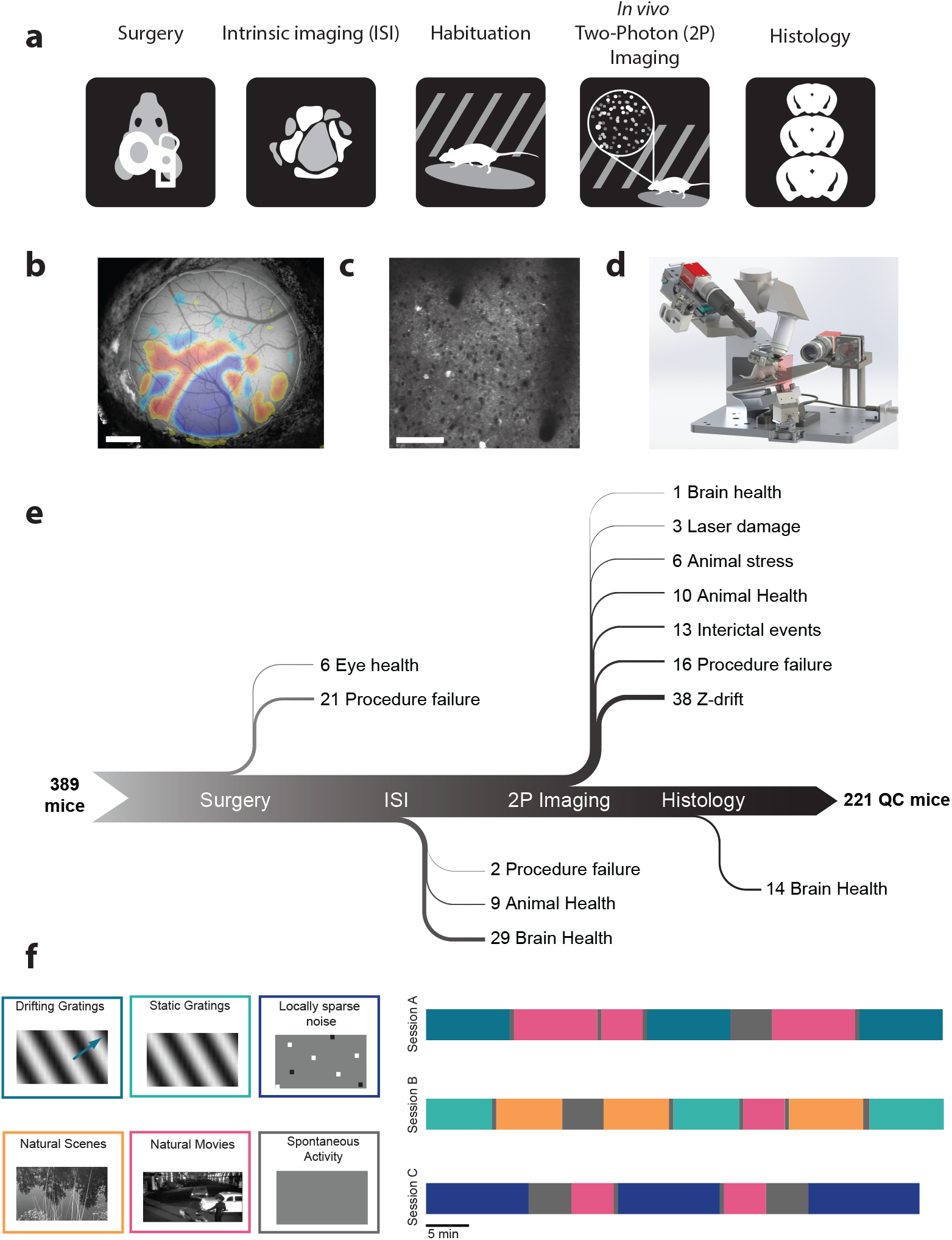
A standardized systems neuroscience data pipeline to map visual responses. (a) Schematic describing the experimental workflow followed by each mouse going through a large scale data pipeline. (b) Example intrinsic imaging map labelling individual visual brain areas. Scale bar = 1mm. (c) Example averaged two photon imaging field of view (400 μm x 400 μm) showcasing neurons labeled with Gcamp6f. Scale bar = 100 μm. (d) Custom design apparatus to standardize the handling of mice in two photon imaging. We engineered all steps of the pipeline to co-register data and tools, enabling reproducible data collection and a standardized experimental process (see Supplementary Figure 1-4). (e) Number of mice passing Quality Control (QC) criteria established by Standardized Operating Procedures (SOPs) at each step of the data collection pipeline with their recorded failure reason. The data collection pipeline is closely monitored to maintain consistently high data quality. (f) Standardized experimental design of sensory visual stimuli to map responses properties of neurons across the visual cortex. 6 blocks of different stimuli were presented to mice (left) and were distributed into 3 separate imaging session called session A, session B and session C (right).

**Table 1:** Visual coding dataset. The number of cells (and experiments) imaged for each Cre line in each cortical visual area. In total, 59,526 cells imaged in 410 experiments in 221 mice are included in this dataset.

| Cre line | Layers | E/I | V1 | LM | AL | PM | AM | RL |
| --- | --- | --- | --- | --- | --- | --- | --- | --- |
| Emx1-IRES-Cre; Camk2a-tTA; Ai93 | 2/3, 4, 5 | E | 3096 (9) | 2098 (8) | 1787 (7) | 835 (4) | 457 (3) | 3011 (9) |
| Slc17a7-IRES2-Cre; Camk2a-tTA; Ai93 | 2/3, 4, 5 | E | 1864 (13) | 1864 (10) | 374 (2) | 1202 (10) | 235 (2) | 322 (3) |
| Cux2-CreERT2; Camk2-tTA; Ai93 | 2/3, 4 | E | 5168 (15) | 3845 (13) | 3037 (12) | 2987 (14) | 1611 (10) | 1370 (9) |
| Rorb-IRES2-Cre; Camk2a-tTA; Ai93 | 4 | E | 2218 (8) | 1191 (6) | 1242 (6) | 593 (6) | 735 (8) | 1757 (6) |
| Scnn1a-Tg3-Cre; Camk2a-tTA; Ai93 | 4 | E | 1873 (9) |  |  |  |  |  |
| Nr5a1-Cre; Camk2a-tTA; Ai93 | 4 | E | 702 (8) | 416 (6) | 172 (4) | 331 (7) | 171 (6) | 1318 (5) |
| Rbp4-Cre_KL100; Camk2a-tTA; Ai93 | 5 | E | 531 (8) | 640 (8) | 490 (7) | 590 (7) | 355 (8) | 136 (5) |
| Fezf2-CreER;Ai148 | 5 | E | 490 (5) | 981 (5) |  |  |  |  |
| Tlx3-Cre_PL56;Ai148 | 5 | E | 1181 (6) | 946 (3) |  |  |  |  |
| Ntsr1-Cre_GN220;Ai148 | 6 | E | 331 (4) | 210 (2) |  | 330 (3) |  |  |
| Sst-IRES-Cre;Ai148 | 4, 5 | I | 449 (18) | 413 (16) | 200 (2) | 608 (17) |  | 46 (2) |
| Vip-IRES-Cre;Ai148 | 2/3, 4 | I | 247 (16) | 280 (15) |  | 320 (15) |  |  |

In order to systematically collect physiological data on this scale, we built data collection and processing pipelines (**Figure 1, Supplemental Figures 1-5**). The data collection workflow progressed from surgical headpost implantation and craniotomy to retinotopic mapping of cortical areas using intrinsic signal imaging, *in vivo* two-photon calcium imaging of neuronal activity, brain fixation, and histology using serial two-photon tomography (**Figure 1a,b,c**). To maximize data standardization across experiments, we developed multiple hardware and software tools to regulate systematic data collection (**Figure 1d**). One of the key components was the development of a registered coordinate system that allowed an animal to move from one data collection step to the next, on different experimental platforms, and maintain the same experimental and brain coordinate geometry (**see Methods, Supplemental Figure 1**). In addition to such hardware instrumentation, formalized standard operating procedures and quality control metrics were crucial for the collection of these data (**Figure 1e**).

Following data collection, movies of fluorescence associated with calcium influx were motion corrected, normalized, and regions of interest (ROIs) were segmented using automated algorithms (**see Methods, Supplemental Figure 9**). Signals from overlapping ROIs were demixed, and contamination from surrounding neuropil was subtracted (**Supplemental Figure 10**). Segmented ROIs were matched across imaging sessions and ROIs were filtered to remove apical dendrites and other processes, with the aim of only including somatic ROIs. For each ROI, events were detected from ΔF/F using an L0 regularized deconvolution algorithm (**see Methods, Supplemental Figure 11**), which deconvolves pointwise events assuming a linear calcium response for each event and penalizes the total number of events included in the trace.

For each neuron, we computed the mean response to each stimulus condition using the detected events, and parameterized its tuning properties. Many neurons showed robust responses, exhibiting orientation-selective responses to gratings, localized spatial receptive fields, and reliable responses to natural scenes and movies (**Figure 2a-f**, **Supplemental Figure 13**). For each neuron and each categorical stimulus (i.e. drifting gratings, static gratings, and natural scenes), the preferred stimulus condition was identified as the condition that evoked the largest mean response for that stimulus (e.g. the orientation and temporal frequency with the largest mean response for drifting gratings). For each trial of the stimulus, the neural activity of the neuron was compared to a distribution of activity for that neuron taken during the epoch of spontaneous activity, and a p-value was computed. If at least 25% of the trials of the neuron’s preferred condition had a significant difference from the distribution of spontaneous activities (p<0.05), the neuron was deemed to be responsive to that stimulus (see **Methods** for responsiveness criteria for locally sparse noise and natural movies).

**Figure 2:**
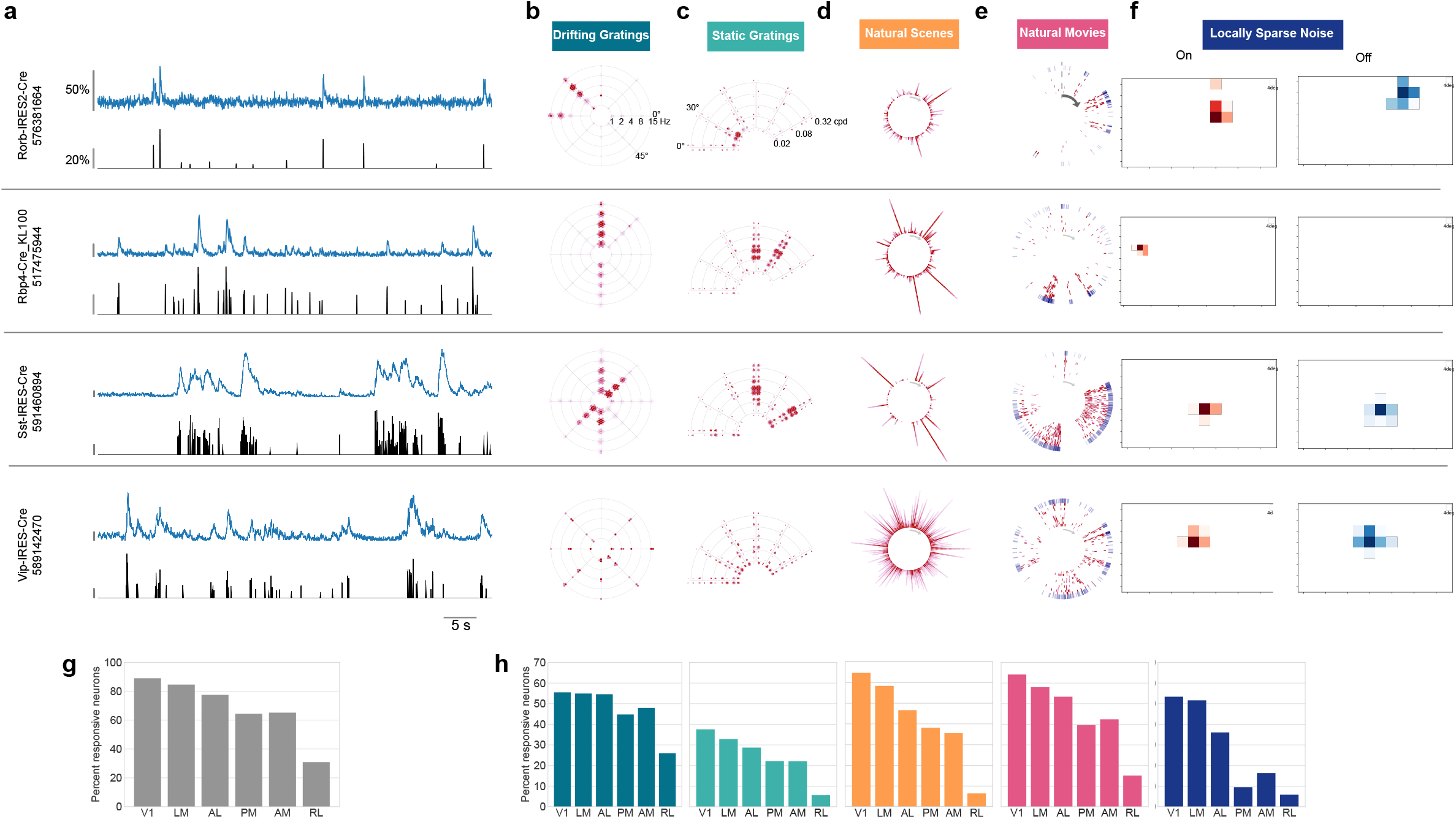
Visual responses to diverse visual stimuli. (a) Activity for four example neurons, two excitatory neurons (Rorb, layer 4, Rbp4, layer 5) and two inhibitory neurons (Sst layer 4, and Vip layer 2/3). ΔF/F (top, blue) and extracted events (bottom, black) for each cell. (b) Star plot summarizing orientation and temporal frequency tuning for responses to the drifting gratings stimulus (For details on response visualizations see Supplemental Figure 13). (c) Fan plot summarizing orientation and spatial frequency tuning for responses to static gratings. (d) Corona plot summarizing responses to natural scenes. (e) Track plot summarizing responses to natural movies. (f) Receptive field subunits mapped using locally sparse noise. (g) Percent of neurons that responded to at least one stimulus across cortical areas. (h) Percent of neurons that responded to each stimulus across cortical areas. Colors correspond to the labels at the top of the figure.

In total, 74% of neurons were responsive to at least one of the visual stimuli presented (**Figure 2g**). The percent of responsive neurons depended on area and stimulus, such that V1 and LM showed the highest number of visually responsive neurons. This dropped in other higher visual areas and was lowest in RL where only 31% of neurons responded to any of the visual stimuli. Natural movies elicited responses from the most neurons, while static gratings elicited responses from the fewest (**Figure 2h**). In addition to varying by area, the percent of responsive neurons was also specific to Cre lines and layers, suggesting functional differences across these dimensions (**Supplemental Figures 14-18**). Note that the retinotopic location of the center of gaze is close the border of RL and somatosensory cortex, which could result in the imaging of non-visual neurons and cause the low rate of responsiveness in this area.

For responsive neurons, visual responses were parameterized by computing several metrics, including preferred spatial frequency, preferred temporal frequency, direction selectivity, and receptive field size (**see Methods**). Comparing these metrics across these areas, layers, and Cre lines, we find evidence of functional differences across these dimensions (**Figure 3**, **Supplemental Figures 19, 20**).

**Figure 3:**
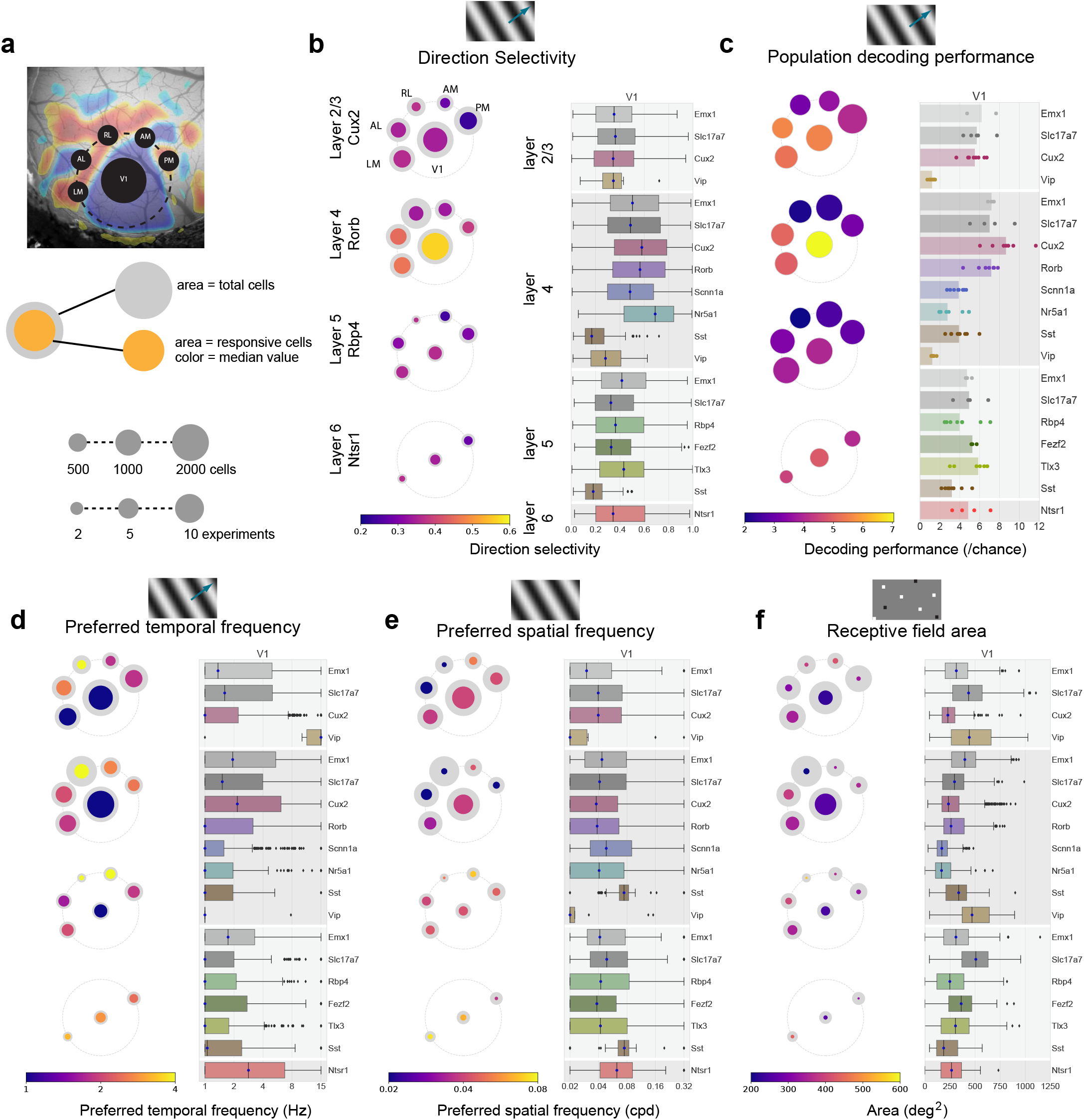
Tuning properties reveal functional differences across areas and Cre lines. (a) Pawplot visualization summarizes median value of a tuning metric across visual areas. Top, each visual area is represented as a circle, with V1 in the center and the higher visual areas surrounding it according to their location on the surface of the cortex. Center, each paw-pad (visual area) has two concentric circles. The area of the larger circle represents the number of cells imaged at that layer and area. The area of the inner, colored, circle represents the number of responsive cells for that layer and area. The color of the inner circle reflects the median value of the metric for the responsive cells, indicated by the colorscale at the bottom of the plot. Bottom, scale of circle area for single cell metrics and for population metrics. In contrast to single-cell metrics, for population metrics (e.g. Fig 3c) each paw-pad (visual area) has only one circle, and the area represents the number of datasets. For a metric’s summary plot, four pawplots are shown, one for each layer. Only data from one Cre line is shown for each layer. For each panel, a pawplot is paired with a box plot or a strip plot (for single cell and population metrics respectively) showing the full distribution for each Cre line and layer in V1. Data is assigned to cortical layers based on both the Cre line and the imaging depth. Data collected above 275um from the surface is considered to be in layer 2/3. Data collected between 275μm and 375μm is considered to be in layer 4. Data collected between 375μm and 500μm is considered to be in layer 5. Data collected at 550μm in considered to be in layer 6. The box shows the quartiles of the data, and the whiskers extend to 1.5 times the interquartile range. Points outside this range are shown as outliers. For other cortical areas, see Supplemental Figure 19. (b) Pawplot and box plot summarizing direction selectivity. (c) Pawplot and strip plot summarizing decoding performance for drifting grating direction using K-nearest neighbors. Each dot represents the mean five-fold cross-validated decoding performance of a single experiment, with the median performance for a Cre-line/layer represented by bar. (d) Pawplot and box plot summarizing preferred temporal frequencies. (e) Pawplot and box plot summarizing preferred spatial frequencies. (f) Pawplot and box plot summarizing receptive field area.

We included several Cre lines that label specific sub-populations of neurons. For instance, Rorb, Scnn1a-Tg3, and Nr5a1 label distinct layer 4 populations, and exhibit distinct tuning properties. For all the computed parameters, Rorb and Scnn1a-Tg3 show significant differences (KS test, **Supplemental Figure 20**) suggesting distinct channels of feedforward information. In layer 5, on the other hand, Tlx3 and Fezf2, which label cortico-cortico and cortico-thalamic projecting neurons respectively, do not show significant differences, implying more homogenous feedback signals. These data also provide the first broad survey of visually evoked responses of both Vip and Sst inhibitory neurons. Responses to drifting gratings support the model of mutual inhibition between these inhibitory populations^13,14^, wherein nearly all Sst cells respond reliably to the grating stimulus while the Vip cells are nearly all unresponsive, and possibly even suppressed (**Supplemental Figure 14**). Interestingly, receptive fields mapped using locally sparse noise reveal that Vip neurons have remarkably large receptive field areas, larger than both Sst and excitatory neurons in V1 (**Figure 3f**). The visual responses of these two populations add important details to the inhibitory cortical circuit.

Comparisons across areas and layers reveal that direction selectivity is highest in layer 4 of V1 (**Figure 3b**). In superficial layers, the differences across areas indicate that V1, LM, and AL show significantly higher direction selectivity than PM, AM, and RL (**Supplemental Figure 19**). This pattern in single neuron selectivity was reflected in our ability to decode the visual stimulus from single-trial population vector responses, using all cells, responsive and unresponsive. We used a K-nearest-neighbors classifier to predict the grating direction. Matching the tuning properties, areas V1, AL, and LM showed higher decoding performance than AM, PM, and RL, and these differences were more pronounced in superficial layers than in deeper layer (**Figure 3c**). However, there are cases where this relationship between population decoding and direction selectivity is broken. For example, Nr5a1 neurons in V1 show the highest median direction selectivity, but the lowest population decoding performance of excitatory neurons. Even matching population size, Nr5a1 continues to show lower decoding performance than other Cre lines (**Supplemental Figure 21**). Destroying trial-to-trial correlations by shuffling trials, we found a slight increase in decoding performance, indicating that noise correlations do not improve the discriminability of population responses to different stimuli (**Supplemental Figure 21**). This result is in contrast to the impact of noise correlations on population coding in the mammalian retina^15,16^, suggesting a transformation of population coding strategies across the visual pathway.

Across all areas, layers, and stimuli, visual responses in mouse cortex were highly sparse. Among responses to natural scenes, we found that most neurons responded to a very small number of scenes. The sparseness of individual neurons was measured using lifetime sparseness, which captures the selectivity of a neuron’s mean response to different stimulus conditions^17,18^ (**see Methods**). A neuron that responds strongly to only a few scenes will have a lifetime sparseness close to 1, whereas a neuron that responds broadly to many scenes will have a lower lifetime sparseness (**Figure 4a**). Excitatory neurons had a median lifetime sparseness of 0.71 in response to natural scenes. While Sst neurons were comparable to excitatory neurons (median 0.78), Vip neurons exhibited low selectivity (median 0.35). Lifetime sparseness did not increase outside of V1; Responses did not become more selective in the higher visual areas. (**Figure 4b, Supplemental Figures 22,23**). Lifetime sparseness is high for all stimuli (data not shown). A complement to the sparseness of an individual neuron is the population sparseness - a measurement of how many neurons respond to each stimulus condition. Like lifetime sparseness, population sparseness is also high in these data for excitatory and Sst neurons (**Figure 4c**), across all areas.

**Figure 4:**
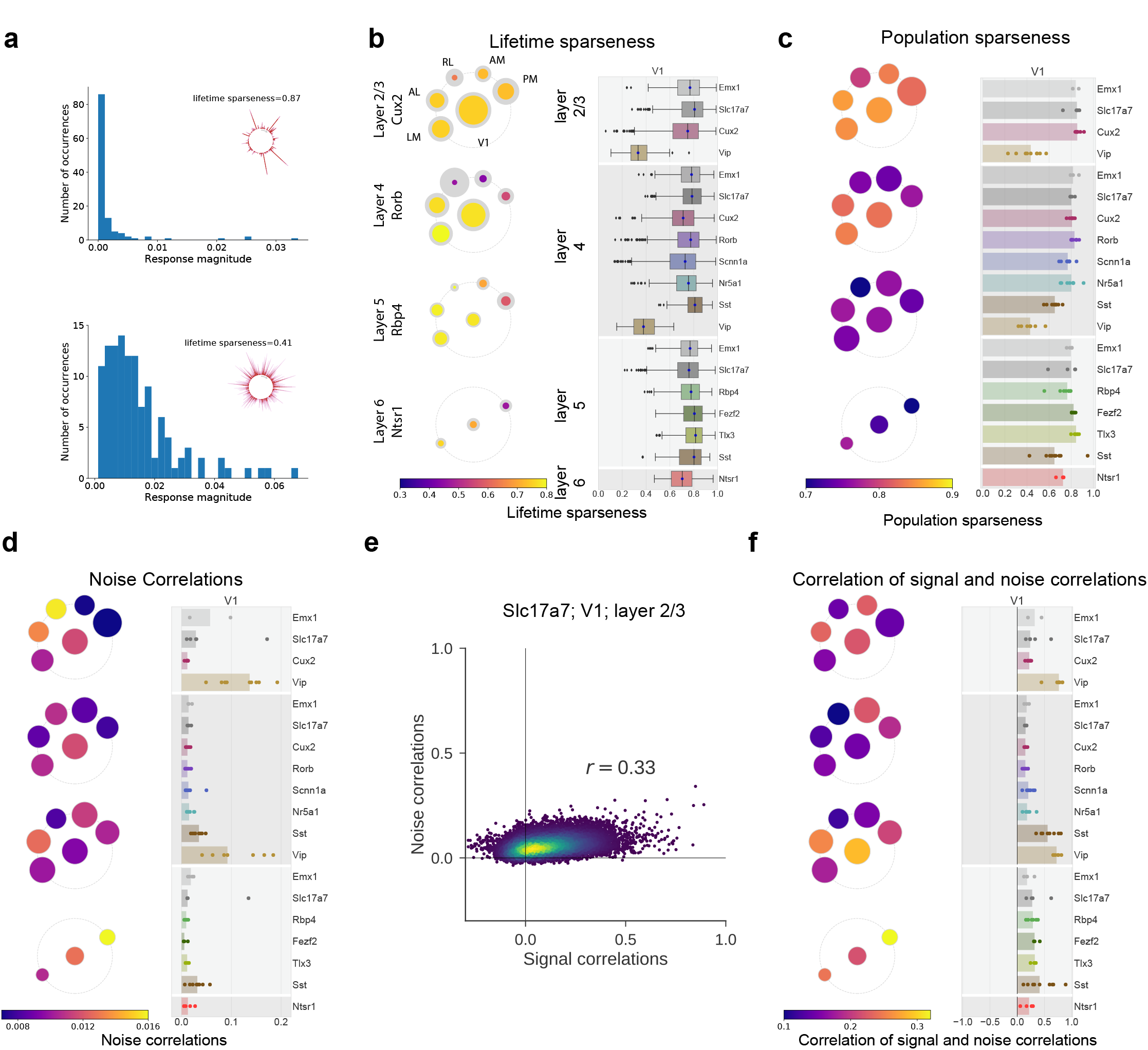
Visual responses are sparse, but coding is dense. (a) Distribution of evoked responses for two example cells showing either high lifetime sparseness (top) and low lifetime sparseness (bottom). The corona plot for each cell is inset in the plot. (b) Pawplot and box plots summarizing lifetime sparseness of the responses to natural scenes. (c) Pawplot and strip plot summarizing the population sparseness of responses to natural scenes. (d) Pawplot and strip plot summarizing the mean noise correlation of responses to natural scenes. (e) Correlation (spearman’s rho) between noise correlations and signal correlations for one experiment (Slc17a7, layer 2/3 of V1). (f) Pawplot and strip plot summarizing the correlation of signal correlations and noise correlations.

Such sparse activity could underlie a form of sparse coding to reduce redundancy and increase efficiency, such that neurons with similar tuning preferences do not respond at the same time.^19,20^ This makes a specific prediction: Similarly tuned neurons should have negatively correlated trial-by-trial activity. Contrary to this prediction of “ explaining away,” we found that similarly tuned neurons exhibited positively correlated trial-by-trial fluctuations in almost all experiments in this dataset (**Figure 4e,f, Supplemental Figure 24**). This result is consistent with reports in other sensory systems and recording methods,^21^ suggesting that sparse single-neuron responses underlying dense population codes are a common feature of cortical representations at the level of rates.

In addition to sparsity in responses across stimulus conditions, the visually evoked responses throughout the mouse cortex showed a large amount of trial-to-trial variability. Indeed, the percent of responsive trials for most neurons at their preferred conditions was low — the median is less than 50% (**Figure 5a, Supplemental Figure 25**). This means that the majority of neurons in the mouse visual cortex are usually unresponsive, even when presented with the stimulus condition that elicits their largest average response. We also calculated a more complete measurement of response reliability, defined as the square of the expected correlation between the trial averaged response to the true, unmeasured, mean response^22^ (**see Methods**). A neuron that responds precisely the same way on each trial to a set of stimuli will have a reliability of 1, while a completely random neuron will have a reliability of 0. We find that neurons had higher reliability for natural stimuli than for the artificial stimuli across all areas and layers (**Figure 5b,c**, Supplemental Figure 25). Altogether, responsive neurons had a mean reliability of 0.62 ± 0.2 (st. dev) for natural scenes and 0.46 ± 0.2 (st. dev) for drifting gratings.

**Figure 5:**
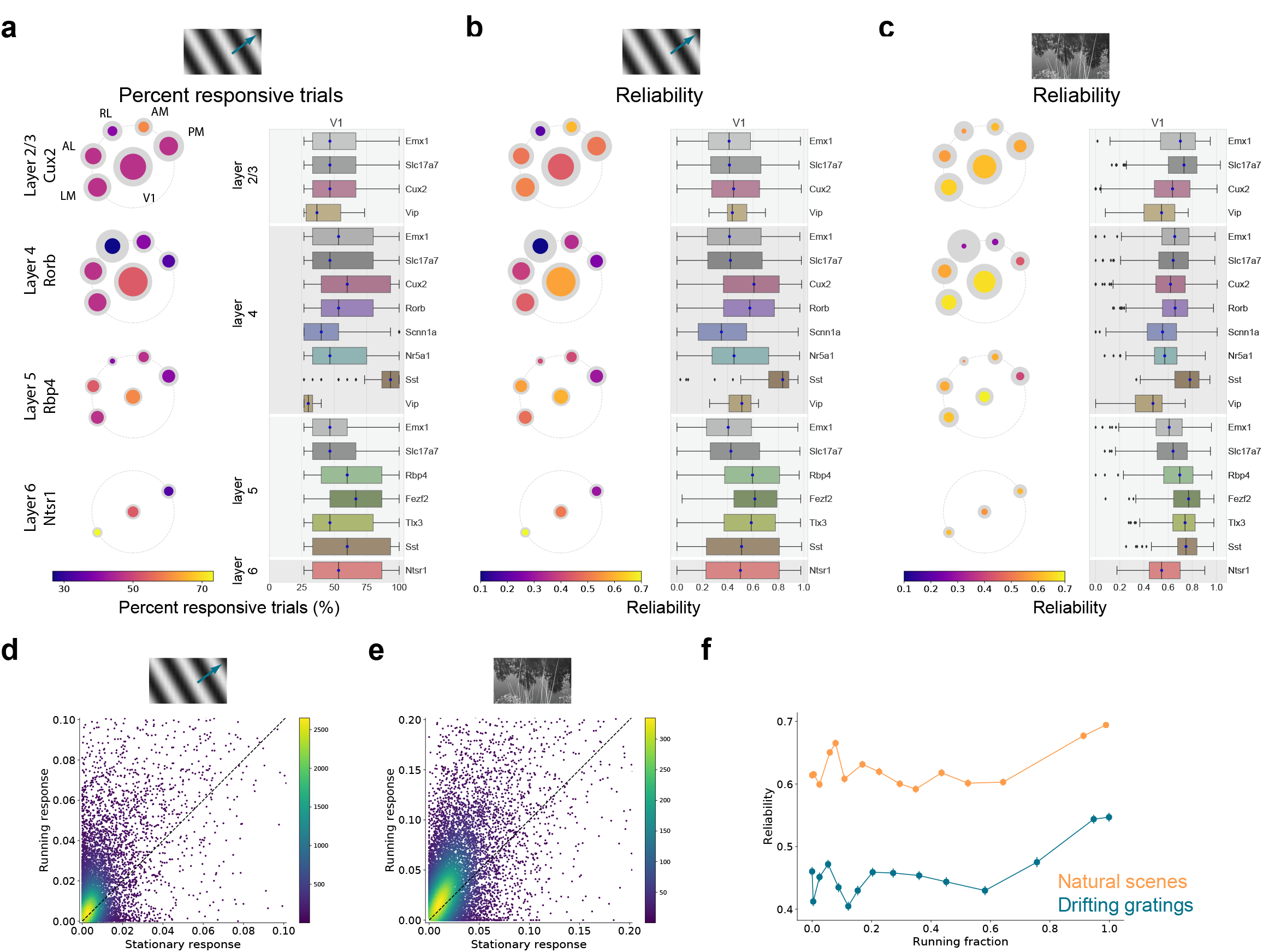
Neural variability is only weakly explained by locomotor activity. (a) Paw plot and box plot summarizing the percent of responsive trials for drifting gratings, the percent of trials that have a significant response for each neurons preferred grating condition. The responsiveness criteria is that a neuron responded to 25% of the trials, hence the low end is capped at 25%. (b) Paw plot and box plot summarizing the reliability of responses for drifting gratings. (c) Paw plot and box plot summarizing the reliability of responses for natural scenes. (d) Evoked response to a neuron’s preferred drifting grating condition when the mouse is running (running speed > 1 cm/s) compared to when it is stationary, shown as a density plot. (e) Evoked response to a neuron’s preferred natural scene when the mouse is running (running speed > 1 cm/s) compared to when it is stationary, shown as a density plot. (f) Reliability as a function of running fraction, data binned into equally sized bins, for drifting gratings and natural scenes.

One possible source of trial-to-trial variability could be the locomotor activity of the mouse. Previous studies have shown that visual responses in the mouse cortex are modulated by the animal’s running activity.^23–27^ The mice in our experiments were free to run on a disc during the experiment and animals showed a range of running behaviors (**Supplemental Figure 26**). For experiments in which the animals spent enough time running such that there were sufficient stimulus trials when the mouse was both stationary and running (at least 10% of trials for each), we compared the responses in these two states. Consistent with previous reports, many neurons show modulated response (**Figure 5d,e**). While most neurons show enhanced responses when running, for many neurons the difference between stationary and running is not significant (only 13% and 37% of neurons show significant modulation of their responses to drifting gratings and natural scenes respectively, using a KS test).

To examine whether the locomotor activity could be a source of trial-to-trial variability, we compared the reliability of neurons’ visual responses to the fraction of time the animal spent running. We found that reliability is higher when the mouse runs consistently, but this increase is modest from a baseline of reliability when the mouse is either completely stationary or shows mixed running behavior (**Figure 5f**). This effect on stimulus response reliability is consistent across different stimuli, both natural and artificial. Thus locomotor activity does contribute to the variability of visual responses, but is unlikely to fully explain the amount of variability found in these data.

We asked whether a standard modeling approach could capture the observed stimulus responses and variability. We used a generalized linear model (GLM) to predict extracted events, smoothed with a Gaussian window, from time series input of the stimuli along with the binary running state of the mouse (**Figure 6a, see Methods**). Only neurons that were matched in all three imaging sessions were used for modeling (∼19,000 neurons), and all neurons were modeled regardless of whether they met our responsiveness criteria. The basis functions for the GLM are two spatiotemporal wavelet pyramids: one a standard linear basis and another that squares the basis functions before summation, approximating a “complex” neuron receptive field. While the model captures the activity of some neurons very well (**Figure 6b**), the median prediction, r, for natural stimuli is ∼0.2-0.3 across areas (**Figure 6c,d**), suggesting a large amount of variation unaccounted for by the stimulus with this model. We computed a complexity ratio by comparing the total weight of the quadratic basis functions to the total weights for each model, and found that almost all neurons are mostly complex, with complexity ratios near 1 (**Figure 6e**). This means that no neuron is better described by a “ simple” linear-nonlinear model than the “complex” quadratic model.

**Figure 6:**
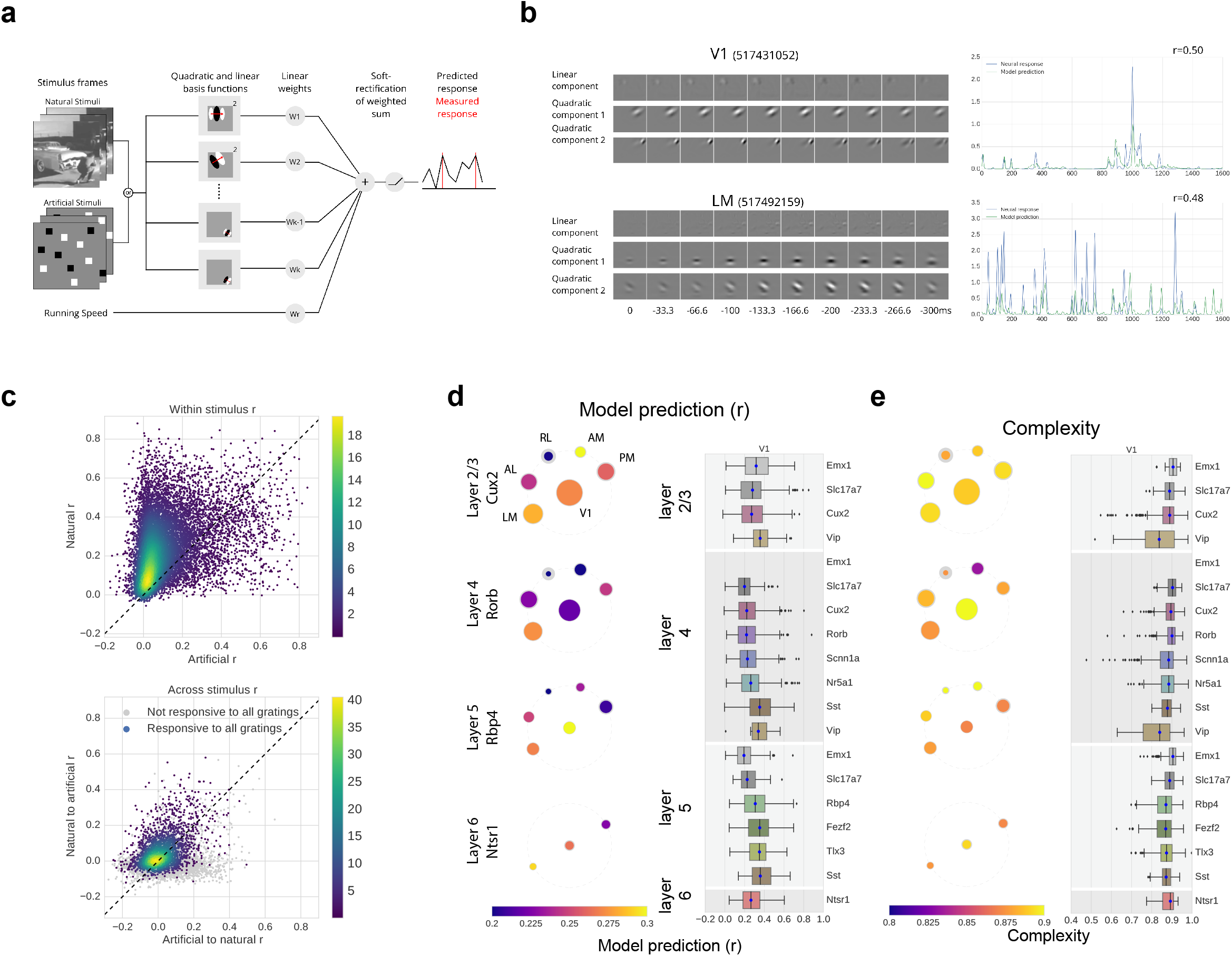
All cells show a high degree of complexity and are better fit with natural stimuli. (a) Schematic for the GLM. The models are trained on either natural or artificial stimuli, converted into a 30Hz time series and spatially downsampled. The time series input is filtered with spatio-temporal Gabor wavelet pyramids, one of which is linearly combined, the other of which is squared before components are combined. These weighted sums are passed through a soft-rectification to predict the detected calcium events, which have been smoothed with a Gaussian filter. (b) (left) Example filters for two cells from the dataset, showing the linear filter as well as two (of many) quadratic components. (right) Predicted response compared with smoothed calcium events for those example cells. (c) Density plot comparing the mean r values for models trained on natural vs. artificial stimuli for all modeled cells (top). Density plot showing cross stimulus performance of models trained on one stimulus type and tested on the other (bottom). (d) Pawplot and box plot summarizing the r values for the dataset. (e) Pawplot and box plot summarizing the complexity across the dataset.

For each neuron, we trained the model separately using natural stimuli (natural scenes and natural movies) and artificial stimuli (drifting gratings, static gratings and locally sparse noise). Comparing the models’ performances, we found that the overall distribution of performance for models trained and tested with natural stimuli was much higher than the corresponding models for artificial stimuli (**Figure 6c**). This was true even for neurons that met our responsiveness criteria for gratings but not natural scenes. Further, models trained on natural stimuli predicted responses to artificial stimuli better than vice versa, although the cross-stimulus prediction was worse than the within-stimulus prediction, consistent with previous reports^9–11^.

Surprisingly, whether a neuron responded to one stimulus (e.g. natural scenes, drifting gratings, etc.) was largely, though not completely, independent of whether it responded to another stimulus. Unlike the examples shown in Figure 2, which were chosen to highlight responses to all stimuli, most neurons were responsive to only a subset of the stimuli presented (**Figure 7a**). The overlap of the set of neurons that responds to each pairwise combination of stimuli was computed for each experiment and compared to the maximum and minimum amount of overlap possible given the fraction of responsive neurons to each stimulus (**Figure 7b, Supplemental Figure 28**). There is above chance overlap for all presentations of natural movies — particularly for natural movie one, whichis repeated in each imaging session (**Figure 7c**). There is also above chance overlap for responses to static gratings and natural scenes. However, natural movies and all other stimuli showed overlap close to the level of chance. That is, whether a neuron responded to natural scenes is independent of whether it responded to natural movies. Notably, locally sparse noise showed the least amount of overlap with other stimuli, and even below chance overlap with some, such as static gratings. These results are consistent across all visual areas.

**Figure 7:**
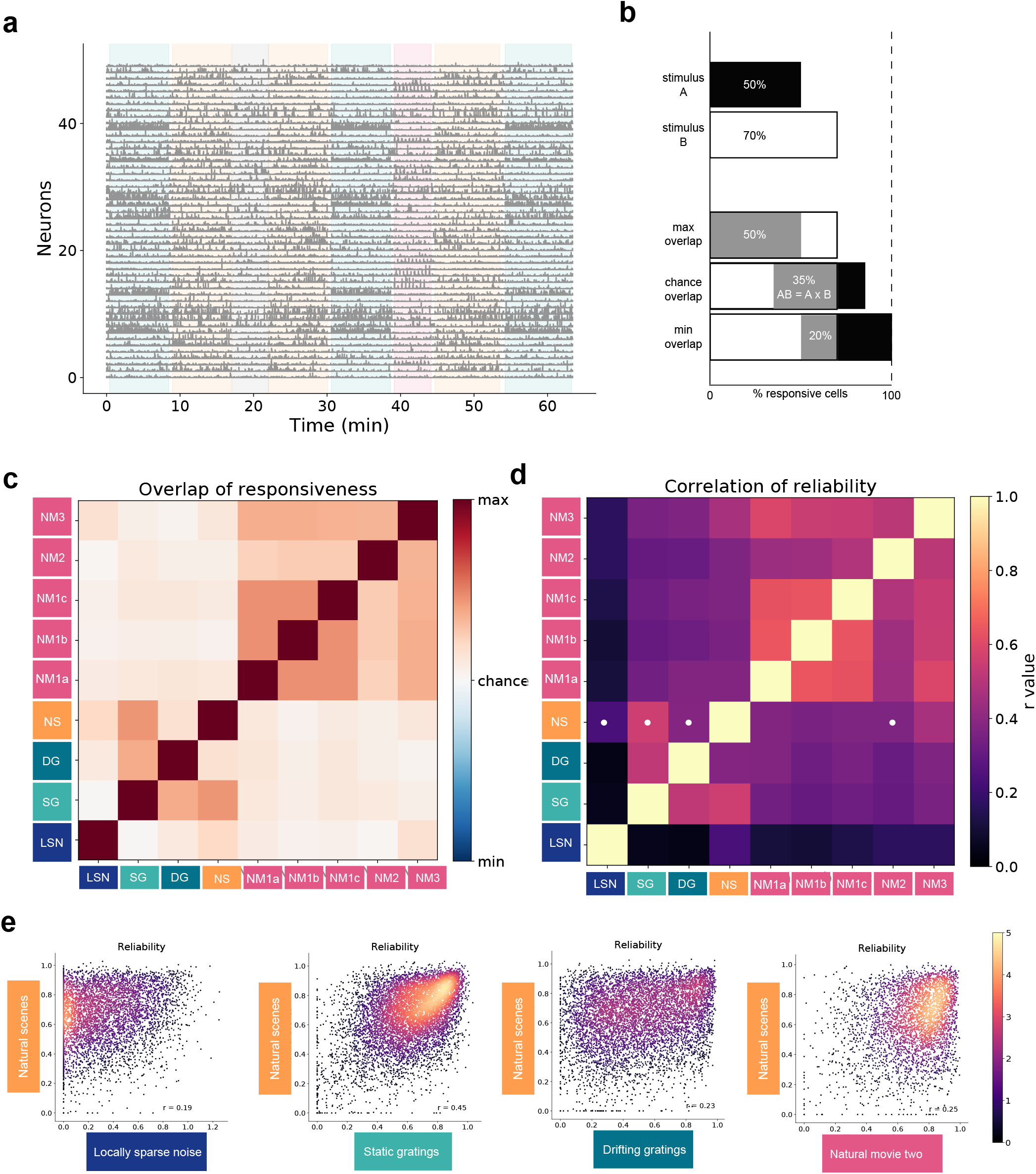
Responses to different stimuli are largely independent. (a) Responses of 50 neurons during one imaging session (Cux2, V1, layer 2/3) with stimulus epochs shaded using stimulus colors from Figure 1. (b) Schematic of overlap analysis. If 50% of cells in an experiment respond to stimulus A and 70% of the cells response to stimulus B, chance overlap would be 35%. Maximum overlap would be 50%, and minimum overlap would be 20%. The overlap between each pair of stimuli was computed, and z-scored. (c) Median overlap z-score for each pair of stimuli for all experiments. (d) The correlation of response reliability for cells responses to each pair of stimuli. White dots indicate the combinations that are shown in panel d (e) Comparison of the reliability of responses for natural scenes with locally sparse noise, static gratings, drifting gratings and natural movie three (left to right).

The independence of whether a neuron responded to two stimuli is also reflected in the correlation between the reliability of neurons’ responses to those two stimuli (**Figure 7d,e**). For neurons that responded to two stimuli, we computed the Pearson correlation between the reliability of responses to each stimulus. We found the same structure in cross-stimulus comparisons such that the reliabilities of natural movie responses were highly correlated, but most stimulus pairs had low correlations. Thus, whether a neuron responds to two stimuli is largely independent, and even when it does respond to both, the reliability of those responses remains largely independent.

Independence between responses to ostensibly similar stimuli is a striking feature of the data and one not predicted by the classical model of the early visual system (namely spatiotemporal Gabor-type wavelets). This observation, together with the fact that neural activity is sparse in both a lifetime and population sense, and finally that the “ simple” linear-nonlinear wavelet based GLM accounted for so little of the explainable variance, all point to the idea that much of the neural activity is driven by relatively higher order features. We quantified this by comparing the population level neural responses to standard deep convolutional networks (CNNs; **Figure 8**). This is an interesting comparison because the original inspiration for these model architectures was the important and early set of results describing “ simple” and “ complex” neurons in Area 17 of anesthetized cat visual cortex^1^.

**Figure 8:**
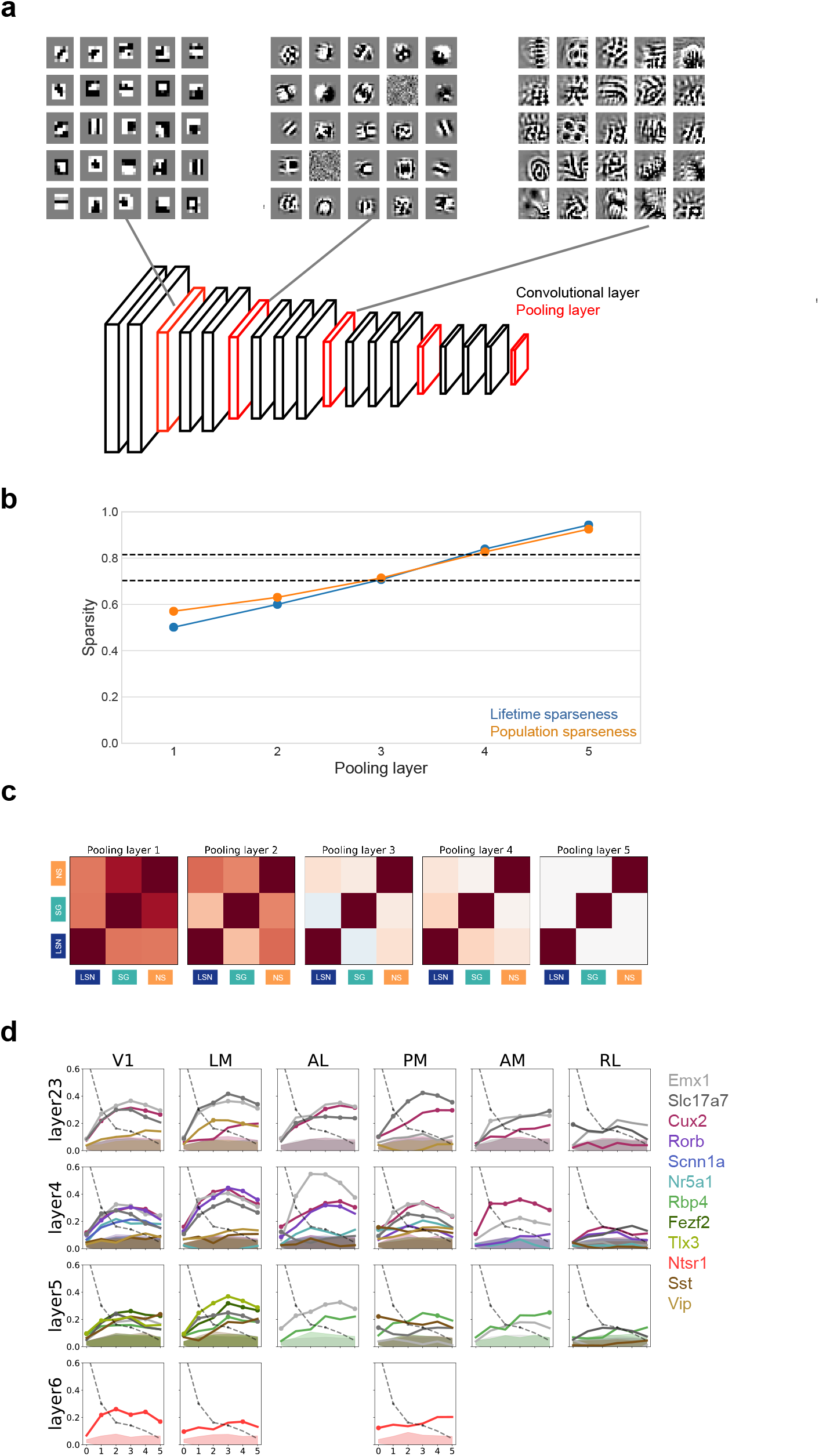
Mouse visual cortex maps to mid-to-high levels of a Convolutional Neural Network. (a) Schematic of VGG16 showing convolutional (black) and pooling (red) layers. Above, example optimal stimuli for sample units found at the first three pooling layers. (b) Median lifetime and population sparseness for each pooling layer of VGG16 in response to the natural scenes stimulus used for this dataset. Dashed lines indicate the limits of median lifetime sparseness for natural scenes found in V1 (see Figure 4b). (c) Stimulus overlap (for the flashed stimuli from the data set) for the pooling layers of VGG16. (d) Similarity of similarity matrix correlation between neural data from each Cre line, area, layer and each pooling layer of VGG16 (see Methods). Shaded region is the null distribution for significance at one standard deviation. Dashed line indicates the SSM correlation with a spatial wavelet pyramid.

Units in CNN models (such as VGG16^28^) are optimally driven by progressively higher order features in deeper layers of the model (**Figure 8a**). The first pooling layer contains many units that appear as coarse edge detectors, while the second pooling layer contains more complex features, with a small subset consisting of oriented gratings similar to traditional V1 receptive fields. By the third pooling layer, there are no such simple looking features, but even more complex shapes and textured patterns. As a natural consequence of this increasing specificity, we see the lifetime and population sparsity in response to natural scenes increase through the pooling layers (**Figure 8b**). This trend is consistent across multiple CNNs; It is not specific to VGG16 (not shown).

Units in VGG16 also display the independence of stimuli observed in the data (**Figure 8c**). We compared the units that respond to each of the flashed stimuli (locally sparse noise, static gratings and natural scenes) for each pooling layer of VGG16. For the lower layers, as expected, there is a high degree of overlap in populations that respond to different stimuli. Moving through deeper layers of the network, the degree of independence increases. The last pooling layer shows nearly complete independence of stimuli.

We used similarity-of-similarity matrix (SSM) analysis^29^ to compare the neural responses with responses at different pooling layers of VGG16 in order to quantify how similar the two representations are (**Figure 8d**). A similarity matrix is constructed by computing the correlation of the trials average population responses to pairs of scenes. We then computed the correlation of similarity matrices between each pooling layer of VGG16 and each cortical area, layer and Cre line in these data. Because the network has a degree of similarity to itself, we only compare pooling layers as the model layers between pooling layers are highly correlated (see **Methods**).

The highest correlations are for pooling layer 3 of VGG16 for most cortical areas and layers (**Figure 8d**). Superficial layers in V1 map to the middle layers most strongly whereas LM, PM, and AL in those layers tend to map to slightly higher layers, suggesting a potential hierarchy, albeit a shallow one^30^. As a comparative baseline, we compute the SSM metric for a linear Gabor wavelet basis (**Figure 8d**), which is highest in the input layer and falls deeper into the network. These results support the view that throughout the mouse visual cortex, neurons exhibit responses to more complex and sophisticated stimuli than the classical model suggests.^5,31^

## Discussion

Data standardization and experimental reproducibility is both a challenge and an opportunity for the field of systems neuroscience. *In vivo* neuronal recordings are notoriously difficult experiments that require an in-depth expertise in many scientific fields and multiple years of training. As such, these experiments are difficult to scale up. Despite these challenges, large cohesive datasets for systems neuroscience offer an opportunity to address fundamental issues of standardization and reproducibility. Here we combined standardized operating procedures with integrated engineering tools to address these long-standing difficulties. We demonstrated data collection in over an order of magnitude more animals (221 mice) than is typically performed in the field and maintained tight standardization across three years of continuous data collection.

We have reduced critical experimental biases by separating quality control of data collection from response characterization. Historically, the field has been dominated by single-neuron electrophysiological recordings in which electrodes were advanced until a neuron was found that responded to a test stimulus. The stimulus was then optimized to elicit the strongest reliable response from that neuron. The experiment proceeded using manipulations around this stimulus condition that had been tuned to drive the strongest response. Such studies have discovered many characteristic response properties, but may fail to capture the variability of responses, the breadth of features that elicit a neural response, and the breadth of features that do not elicit a response. Recently, calcium imaging and denser electrophysiological recordings have enabled large populations of neurons to be recorded simultaneously. By combining calcium imaging with strong quality control and standardization, we have created an unprecedented survey of mouse visual cortex using a standard and well-studied but diverse set of stimuli while limiting the selection bias towards those stimuli.

Under the canonical model, V1 sits at the initial stages of a processing hierarchy where neurons respond to low-level features, specifically with spatially localized receptive fields with spatial and temporal frequency preferences.^1–4^ Neural responses become increasing specialized in the higher areas moving away from V1, reaching extremes in which cells show very selective responses to specific objects and even faces.^4,32^

The field has a growing body of evidence showing that the canonical model needs to be enhanced to support more sophisticated visual computation.^5,31^ For instance, neurons in mouse V1 show complex visual responses previously associated with higher cortical areas, including pattern selectivity for plaid stimuli^33^ Furthermore, the emergence of the rodent as a prominent model of the visual system in recent years has revealed evidence of non-visual computation, including behavioral responses such as reward timing and sequence learning^34^, as well as modulation by multimodal sensory stimuli^35,36^ and motor signals.^23,24,37–39^

We expected this survey to provide strong evidence for low-level responses that become progressively higher order throughout the higher visual areas of mouse cortex. Instead, neurons throughout the mouse cortex show highly variable, sparse responses, best fit by “ complex” models. Further, responsiveness to different stimuli is largely independent. Rather than support the canonical model, these results provide evidence of higher order coding wherein neurons exhibit specialized responses to a set of sparse and higher level features of the visual field.

Neurons tuned to low-level features will not, as a whole, show the property of independence that we observe in these data. Such neurons should be, by and large, equally mappable using noise stimuli, grating stimuli, and natural stimuli – with some stimulus specific modifications in the resulting receptive field.^8,9,11^ While we observe individual examples of neurons that behave exactly this way, this is not a general feature of the population of responses (**Figure 7, Supplemental Figure 28**). Computationally, we can consider how a system that responds to low-order features will behave by examining either the early layers of a CNN (**Figure 8c,d**) or a wavelet basis (not shown), where we see strong dependence and correlation of responses across stimuli, contrary to what is observed in the dataset. Strikingly, the fact that none of the neurons in the dataset are better fit by the “ simple” model in our GLM wavelet basis model (**Figure 6**) further supports our finding that neurons are not tuned to low-level features.

Neurons that respond to higher-order features, on the other hand, result in responses that are sparse in both a population and lifetime sense, as we observe here. In a CNN, the network develops features during training that allow it to correctly classify images. Whereas the early units of these networks tend to be more general and low-order, as described above, the intermediate units become increasingly specialized for features that are necessary for the trained task. As a result, the CNN shows a greater degree of stimulus independence with depth (**Figure 8c**). Our data, throughout the mouse visual cortex, shows a degree of independence that is similar to that observed in the third pooling layer of VGG16 (**Figure 8c**). This is consistent with a comparison of sparsity, both lifetime and population, between the dataset and VGG16, as well as the representation mapping using SSM-analysis that shows most layers and areas are more similar to the middle pooling layers, while a wavelet basis is most similar to the input and early layers (**Figure 8b,d**). These results are also consistent with an alternative methodology, SVCCA^40^ (not shown). Taken together, these results reveal that neurons throughout the mouse visual cortex exhibit higher order coding, revealing that they are specialized for high-level features.

This is not to say that there are not plenty of cells in the early visual cortical areas that show Gabor-type receptive fields. VGG16, at the second pooling layer, for example, has units with optimal stimuli that closely resemble Gabors, but they are the minority. Additionally, probing such networks with stimuli such as linear gratings or noise stimuli, or with approaches such as spike triggered averaging, will result in responses that can be characterized with Gabor-type receptive fields even though this is not the optimal stimulus condition that drives such units. We posit that the same phenomenon is almost certainly at play in the mouse visual cortex. Specialized, higher-order visual neurons have been known to exist, either high in the visual hierarchy or as particular exceptions (e.g. loom detectors, motion pattern cells). By including a broad range of stimuli and reducing stimulus bias in our data collection and analysis, we have revealed that such higher order cells are closer to the rule than the exception in the mouse visual cortex. Given that much of the existing literature describes the visual system of cat and primate, it is interesting to speculate as to whether these results might generalize to other species.

Identifying the exact response characteristics of the population of cells remains an open problem. The optimal stimuli of units in a CNN are the result of optimization for an object recognition task on natural stimuli. Such a “ task” appropriate to define the response characteristics of the mouse visual system remains unclear. Understanding the computation of the mouse visual circuit will require identifying the features and stimuli that are ethologically relevant to the mouse.

The Allen Brain Observatory Visual Coding dataset is an openly available dataset, accessible via a dedicated web portal (http://observatory.brain-map.org/visualcoding), with a custom Python-based Application Programming Interface, the AllenSDK (http://alleninstitute.github.io/AllenSDK/). We believe these data will be a valuable resource to the systems neuroscience community as a testbed for theories of cortical computation and a benchmark for experimental results. Already, these data have been used by other researchers to develop image processing methods,^41,42^ to examine stimulus encoding and decoding,^43–47^ and to test models of cortical computations.^48^ Ultimately, we expect these data will seed as many questions as they answer, fueling others to pursue both new analyses and further experiments to unravel how cortical circuits represent and transform sensory information.

## Acknowledgements

We thank the Animal Care, Transgenic Colony Management and Lab Animal Services for mouse husbandry. We thank Daniel Denman, Josh Siegle, Yazan Billeh and Anton Arkhipov for critical feedback on the manuscript. This work was supported by the Allen Institute, and in part by NIH 2R90DA033461, NSF DMS-1514743, Falconwood Foundation, Center for Brains, Minds & Machines funded by NSF Science and Technology Center Award CCF-1231216, Natural Sciences and Engineering Research Council of Canada, NIH Grant DP5OD009145, and NSF CAREER Award DMS-1252624. We thank Allan Jones for providing the critical environment that enabled our large scale team effort. We thank the Allen Institute founder, Paul G Allen, for his vision, encouragement, and support.

## Author Contributions

SEJdV, MAB, KR, MG, TK, SM, SO, JW, CD, LN, AB, JWP, RCR, and CK conceived of and designed the experiment. JL, TK, PH, AL, CS, DS, and CF built and maintained the hardware. SEJdV, JL, MAB, GKO, DF, NC, LK, WW, DW, RV, CB, BB, TD, JG, SJ, NK, CL, FL, FL, JP, NS, DW, JZ, and LN developed algorithms and software, including the SDK and website. KR, NB, NB, SC, LC, AC, SC, ME, NG, FG, RH, LH, UK, JL, RL, EL, LL, JL, KM, TN, MR, SS, CW, and AW collected data. JL and PAG managed the pipeline. SEJdV, JL, MAB, GKO, MO, NC, PL, DM, and RV analyzed data. SEJdV, JL, and MAB wrote the paper with input from PAG, GKO, MO, NC, PL, DM, RCR, and MG.

## References

1. Hubel, D. & Wiesel, T. Receptive fields of single neurones in the cat’s striate cortex. J. Physiol. 148, 574–591 (1959).

2. Hubel, D. H. & Wiesel, T. N. Receptive fields, binocular interaction and functional architecture in the cat’s visual cortex. J. Physiol. 160, 106–154.2 (1962).

3. Felleman, D. J. & Van Essen, D. C. Distributed Hierarchical Processing in the Primate Cerebral Cortex.

4. DiCarlo, J. J., Zoccolan, D. & Rust, N. C. How does the brain solve visual object recognition? Neuron (2012). doi:10.1016/j.neuron.2012.01.010

5. Olshausen, B. & Field, D. What is the other 85 % of V1 doing? Probl. Syst. &1–29 (2004). doi:10.1093/acprof:oso/9780195148220.003.0010

6. Andermann, M. L., Kerlin, A. M., Roumis, D. K., Glickfeld, L. L. & Reid, R. C. Functional specialization of mouse higher visual cortical areas. Neuron 72, 1025–1039 (2011).

7. Marshel, J. H., Garrett, M. E., Nauhaus, I. & Callaway, E. M. Functional specialization of seven mouse visual cortical areas. Neuron 72, 1040–1054 (2011).

8. Fournier, J., Monier, C., Pananceau, M. & Frégnac, Y. Adaptation of the simple or complex nature of V1 receptive fields to visual statistics. Nat. Neurosci. 14, 1053–60 (2011).

9. David, S., Vinje, W. & Gallant, J. L. Natural Stimulus Statistics Alter the Receptive Field Structure of V1 Neurons. J. Neurosci. 24, 6991–7006 (2004).

10. Talebi, V. & Baker, C. L. Natural versus Synthetic Stimuli for Estimating Receptive Field Models: A Comparison of Predictive Robustness. J. Neurosci. 32, 1560–1576 (2012).

11. Yeh, C.-I., Xing, D., Williams, P. & Shapley, R. Stimulus ensemble and cortical layer determine V1 spatial receptive fields. Proc. Natl. Acad. Sci. 106, 14652–14657 (2009).

12. Averbeck, B. B., Latham, P. E. & Pouget, A. Neural correlations, population coding and computation. Nature Reviews Neuroscience(2006). doi:10.1038/nrn1888

13. Pfeffer, C. K., Xue, M., He, M., Huang, Z. J. & Scanziani, M. Inhibition of inhibition in visual cortex: the logic of connections between molecularly distinct interneurons. Nat. Neurosci. 16, 1068–1076 (2013).

14. Fu, Y. et al. A cortical circuit for gain control by behavioral state. Cell 156, 1139–1152 (2014).

15. Franke, F. et al. Structures of Neural Correlation and How They Favor Coding. Neuron (2016). doi:10.1016/j.neuron.2015.12.037

16. Zylberberg, J., Cafaro, J., Turner, M. H., Shea-Brown, E. & Rieke, F. Direction-Selective Circuits Shape Noise to Ensure a Precise Population Code. Neuron (2016). doi:10.1016/j.neuron.2015.11.019

17. Rolls, E. T. & Tovee, M. J. Sparseness of the neuronal representation of stimuli in the primate temporal visual cortex. J. Neurophysiol. 73, 713–726 (1995).

18. Vinje, W. E. & Gallant, J. L. Sparse Coding and Decorrelation in Primary Visual Cortex During Natural Vision. Science (80-.). 287, 1273–1276 (2000).

19. Olshausen, B. A. & Field, D. J. Sparse coding with an overcomplete basis set: A strategy employed by V1 doing? Vis. Res. 37, 3311–3325 (1997).

20. Barlow, H. Possible principles underlying the transformation of sensory messages. Sens. Commun. 217–234 (1961). at http://www.trin.cam.ac.uk/horacebarlow/21.pdf.

21. Cohen, M. R. & Kohn, A. Measuring and interpreting neuronal correlations. Nature Neuroscience (2011). doi:10.1038/nn.2842

22. Schoppe, O., Harper, N. S., Willmore, B. D. B., King, A. J. & Schnupp, J. W. H. Measuring the Performance of Neural Models. Front. Comput. Neurosci. 10, 1–11 (2016).

23. Niell, C. M. & Stryker, M. P. Modulation of visual responses by behavioral state in mouse visual cortex. Neuron 65, 472–9 (2010).

24. Saleem, A., Ayaz, A., Jeffery, K., Harris, K. & Carandini, M. Integration of visual motion and locomotion in mouse visual cortex. Nat. Neurosci. 16, 1864–1869 (2013).

25. Dadarlat, M. C. & Stryker, M. P. Locomotion enhances neural encoding of visual stimuli in mouse V1. 37, 3764–3775 (2017).

26. Dipoppa, M. et al. Vision and locomotion shape the interactions between neuron types in mouse visual cortex. bioRxiv 058396 (2016). doi:10.1101/058396

27. Polack, P. O., Friedman, J. & Golshani, P. Cellular mechanisms of brain state-dependent gain modulation in visual cortex. Nat. Neurosci. 16, 1331–1339 (2013).

28. Simonyan, K. & Zisserman, A. Very Deep Convolutional Networks for Large-Scale Image Recognition. 1–14 (2014). doi:10.1016/j.infsof.2008.09.005

29. Kriegeskorte, N., Mur, M. & Bandettini, P. Representational similarity analysis - connecting the branches of systems neuroscience. Front. Syst. Neurosci. 2, 4 (2008).

30. Harris, J. A. et al. The organization of intracortical connections by layer and cell class in the mouse brain. bioRxiv 292961 (2018). doi:10.1101/292961

31. Masland, R. H. & Martin, P. R. The unsolved mystery of vision. Curr. Biol. 17, R577–82 (2007).

32. Quiroga, Q., Reddy, L., Kreiman, G., Koch, C. & Fried, I. Invariant visual representation by single neurons in the human brain. Nature 435, 1102–1107 (2005).

33. Palagina, G., Meyer, J. F. & Smirnakis, S. M. Complex Visual Motion Representation in Mouse Area V1. J. Neurosci. 37, 164–183 (2017).

34. Gavornik, J. P. & Bear, M. F. Higher brain functions served by the lowly rodent primary visual cortex. Learn. Mem. 21, 527–533 (2014).

35. Bieler, M. et al.Rate and Temporal Coding Convey Multisensory Information in Primary Sensory Cortices. Eneuro 4, ENEURO.0037–17.2017 (2017).

36. Ibrahim, L. A. et al.Cross-Modality Sharpening of Visual Cortical Processing through Layer-1-Mediated Inhibition and Disinhibition. Neuron 89, 1031–1045 (2016).

37. Keller, G. B., Bonhoeffer, T. & Hübener, M. Sensorimotor mismatch signals in primary visual cortex of the behaving mouse. Neuron 74, 809–15 (2012).

38. Stringer, C. et al.Spontaneous behaviors drive multidimensional, brain-wide population activity. bioRxiv 306019 (2018). doi:10.1101/306019

39. Musall, S., Kaufman, M. T., Gluf, S. & Churchland, A. Movement-related activity dominates cortex during sensory-guided decision making. bioRxiv 308288 (2018). doi:10.1101/308288

40. Raghu, M., Gilmer, J., Yosinski, J. & Sohl-Dickstein, J. SVCCA: Singular Vector Canonical Correlation Analysis for Deep Learning Dynamics and Interpretability. 1–17 (2017). doi:1706.05806

41. Petersen, A., Simon, N. & Witten, D. SCALPEL: Extracting Neurons from Calcium Imaging Data. 1–31 (2017). at http://arxiv.org/abs/1703.06946.

42. Sheintuch, L. et al.Tracking the Same Neurons across Multiple Days in Ca2+Imaging Data. Cell Rep. 21, 1102–1115 (2017).

43. Ellis, R. J. et al.High-accuracy Decoding of Complex Visual Scenes from Neuronal Calcium Responses. 1–32 (2018).

44. Cai, L., Wu, B. & Ji, S. Neuronal Activities in the Mouse Visual Cortex Predict Patterns of Sensory Stimuli. (2018).

45. Zylberberg, J. Untuned but not irrelevant: A role for untuned neurons in sensory information coding. bioRxiv 1–18 (2017). doi:10.1101/134379

46. Christensen, A. J. & Pillow, J. W. Running reduces firing but improves coding in rodent higher-order visual cortex. bioRxiv 1–14 (2017).

47. Esfahany, K., Siergiej, I., Zhao, Y. & Park, I. M. Organization of Neural Population Code in Mouse Visual System. bioRxiv 1–16 (2017). doi:10.1101/220558

48. Sweeney, Y. & Clopath, C. Population coupling predicts the plasticity of stimulus responses in cortical circuits. (2018). doi:10.1101/265041

